# Optimized conditions for GTP loading of Ras

**DOI:** 10.1101/2025.08.08.669329

**Authors:** Kimberly J. Vish, Asha P. Rollins, Maxum E. Paul, Titus J. Boggon

## Abstract

Ras and the small GTPase group are essential for myriad cellular processes and cycle between GDP- and GTP-loaded states to allow stringent control of downstream signaling pathways. Biochemical studies of small GTPases can therefore require a specific nucleotide-bound state. Small GTPases possess basal intrinsic activity to process GTP into GDP plus inorganic phosphate, therefore *in vitro* exchange of GDP for GTP is necessary for assays on GTP-loaded states. Here, we assess the methodology of *in vitro* nucleotide exchange for soluble H-Ras. We begin by describing a protocol to quantify the nucleotide bound content of H-Ras using anion exchange chromatography and use this protocol to investigate optimal strategies for loading Ras with GTP by assessing the effects of time, temperature, H-Ras concentration, magnesium, excess nucleotide, and isoform identity. We continue by considering storage of GTP-loaded H-Ras and present optimal conditions to minimize intrinsic GTP hydrolysis. Finally, we conclude by investigating the nucleotide composition of recombinantly expressed H-Ras encompassing cancer mutations at residues Gly12, Gly13, and Gln61. We therefore describe methodology to quantitatively analyze the nucleotide content of small GTPases and their mutants, and demonstrate conditions to achieve efficient GTP loading of Ras.

## Introduction

Ras proteins are the founding members of the small GTPase group ^1–4^, a family of proteins that alter their conformational state and signaling outcomes depending on the identity of a bound guanine nucleotide, either GDP or GTP ^5–7^. For example, signaling from Ras to MAP kinase cascades is almost exclusively associated with the GTP-bound state of Ras rather than its GDP-bound state ^6, 8^. Due to this nearly binary distinction between the nucleotide-bound states and their signaling outcomes, cycling between these states is highly regulated, with factors known as the Guanine Exchange Factors (GEFs) and GTPase Activating Proteins (GAPs) facilitating fast transitions between GDP and GTP binding (**Figure 1A**). Biochemical and kinetic analyses of the effects of these assistant proteins continue to provide foundational information about their roles and importance for GTPase cascades.

**Figure 1.**
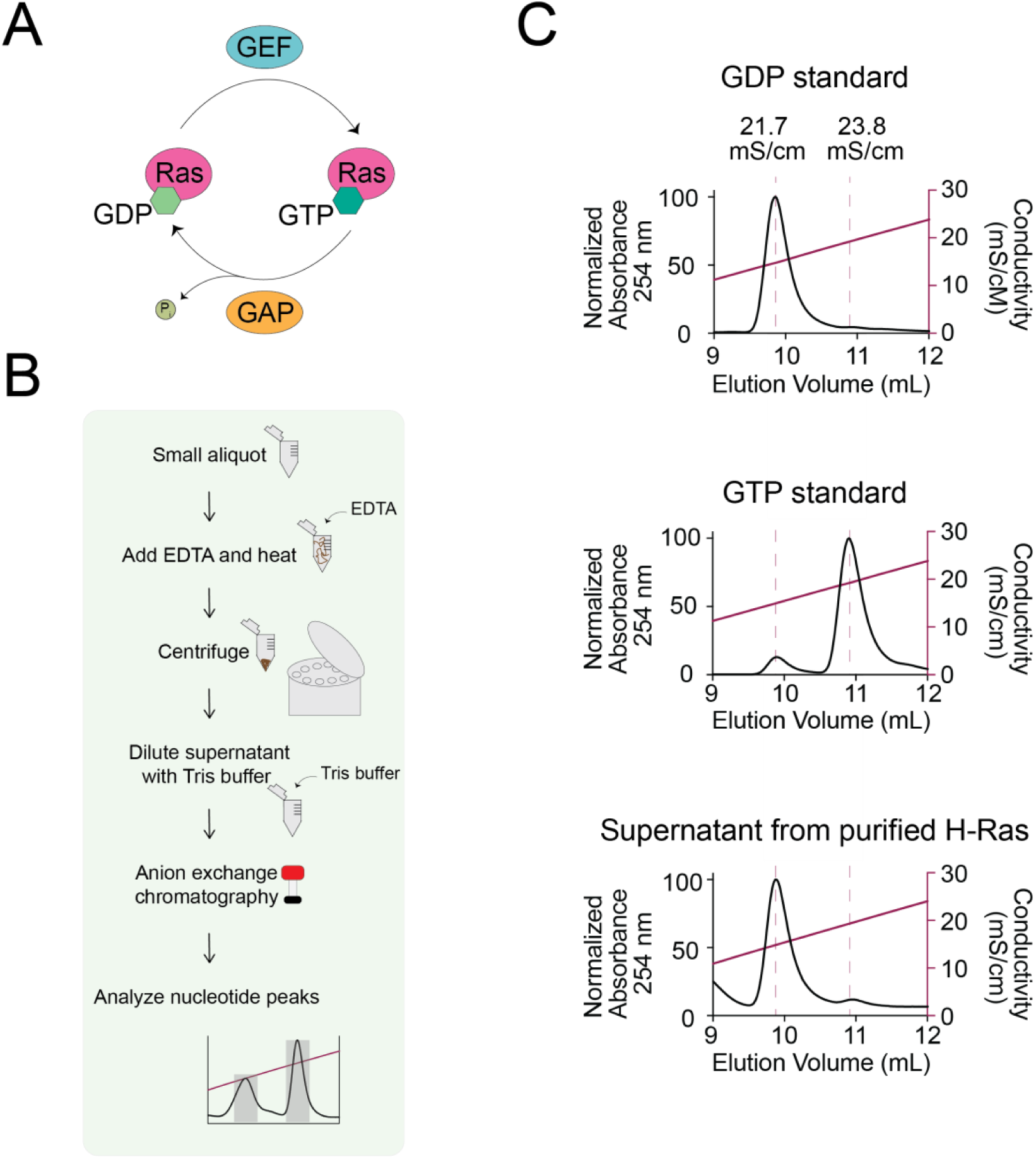
Nucleotide content analysis of GTP-loaded H-Ras. **A)** The GTPase cycle. Nucleotide exchange for small GTPases between the GDP-loaded state and the GTP-loaded state is facilitated by guanine nucleotide exchange factors (GEFs) and the slow intrinsic hydrolysis of GTP to GDP and inorganic phosphate by small GTPases is accelerated by GTPase activating proteins (GAPs). **B)** Flow chart for H-Ras nucleotide content analysis. A small aliquot of purified H-Ras is incubated with EDTA to accelerate nucleotide release, heated to denature the protein and then centrifuged to pellet denatured protein. Supernatant is diluted with Tris buffer to reduce salt concentration to below 50 mM and anion exchange chromatography conducted. Areas under the curve are analyzed for nucleotide peaks to determine ratio of GDP vs GTP bound to H-Ras. **c)** Nucleotide analysis of recombinantly purified H-Ras by anion exchange chromatography. H-Ras purified from recombinant *E. coli* expression was denatured by heating at 95°C for 15 minutes in the presence of 10 mM EDTA, releasing the bound nucleotide to the supernatant. The supernatant was then subjected to anion exchange chromatography and compared to GDP and GTP standards. Peaks are normalized to maximum absorbance.

Ras has four common isoforms arising from three genes, H-Ras, N-Ras, and K-Ras (A and B from alternative splicing arrangements)^9–12^, that have each been extensively studied, particularly because of the high mutational load in cancer and other diseases ^9, 13–15^. At the molecular level, as with other G proteins, Ras small GTPases bind guanine nucleotides at picomolar affinity in the presence of Mg^2+^ ^16–18^. The tight affinity results in strong preference for guanine nucleotide bound versus unbound Ras, and thus promotes slow nucleotide exchange rates, particularly in recombinant expression systems with a lack of appropriate GEF proteins to catalyze exchange. For wild-type Ras, however, intrinsic GTP hydrolysis to GDP+P_i_ occurs at a rate significant on the timescale of recombinant expression ^19–21^ leading to most recombinantly expressed Ras to be GDP bound. Because of this, analyses that require the GTPase in a GTP-loaded state are subjected to biochemical treatment to facilitate exchange of GDP nucleotide for GTP.

Ethylenediaminetetraacetic acid (EDTA) chelation of divalent magnesium induces dissociation of GDP/Mg^2+^ ^22, 23^, an effect that can be exploited in the presence of GTP to facilitate GTPase cycling from GDP-bound to GTP-bound ^24, 25^. EDTA is therefore used in experiments that require loading of Ras with a desired non-GDP nucleotide or nucleotide analog. These include assessments GAP activity on small GTPases (using a variety of nucleotides: e.g. GTP, radiolabeled GTP, or fluorescently tagged) ^26–35^, single turnover assays ^36, 37^, and structural biology of small GTPases and their binding partners ^38–40^. Over the past decades an array of methods have been used to achieve EDTA-mediated nucleotide exchange, with variations in conditions including differences in reaction time, temperature, concentration, and buffer conditions. Importantly, however, the efficiency of loading has not always been assessed, but can quantitatively impact kinetic analyses. Here, we develop an optimal strategy for loading H-Ras with GTP and demonstrate this strategy is applicable to other Ras isoforms. We present an analysis protocol to assess the nucleotide composition of H-Ras using tools commonly found in the biochemistry laboratory. Next, we provide a method for storage of H-Ras in a GTP-loaded state with minimal intrinsic hydrolysis. Finally, we demonstrate the use of these tools to assess the impact of common cancer-associated mutations, G12V, G13R, and Q61L, on GTP loading. We therefore provide optimized methodology for *in vitro* exchange of bound nucleotide to H-Ras.

## Results

### Considerations for the loading of H-Ras with GTP

We began by establishing a protocol for assessing the identity of nucleotide bound to H-Ras. The protocol denatures and removes H-Ras protein but maintains bound nucleotide in the soluble fraction allowing separation by anion exchange chromatography. As adapted from previous work ^41–43^, following H-Ras purification we remove a small aliquot (at least 0.01 nmol of protein, using more for a higher signal) and add 10 mM EDTA to chelate the Mg^2+^ present, facilitating nucleotide release. Incubation of the sample at 95°C for at least 15 minutes releases bound nucleotide and denatures the protein which is then removed by centrifugation. The supernatant contains nucleotide that had previously been bound to H-Ras and is subjected to anion exchange chromatography where a gradient from 0 M – 1.0 M NaCl over 20 column volumes in a buffer of 20 mM Tris pH 8.0 is sufficient to separate GDP and GTP (**Figure 1B**). Importantly for the generalizability of this protocol, both GDP and GTP can be observed at wavelengths of 254 nm and 280 nm, allowing guanine nucleotides to be assessed using common laboratory equipment. Furthermore, both GTP and GDP have the same extinction coefficient which allows direct comparison of integrated peaks as a proxy for amount. When we apply this protocol to assess the nucleotide content bound to the bacterially expressed and purified G-domain of H-Ras, we find that as previously documented by others ^19, 20^ Ras is almost exclusively GDP-bound (**Figure 1C**).

For our biochemical and biophysical assays we desired GTP-bound Ras, not GDP-bound Ras. We therefore assessed the literature to determine an optimal protocol for nucleotide exchange but found what seemed to be large differences in protocol methods. This prompted us to conduct a more extensive inspection of the literature and assess the variability in methodology used to exchange GDP for GTP in small GTPases. In general, published procedures incubate the target small GTPase in the presence of a buffer containing EDTA and excess GTP, followed by buffer exchange to remove excess nucleotide ^34, 44–47^. However, despite this general procedure, careful analysis reveals wide disparities in published experimental procedures (**Table 1**). We find variations in the time of incubation, reaction temperature, amount of excess nucleotide added during EDTA incubation, in the addition of magnesium chloride after EDTA incubation, and also in the inclusion of EDTA or magnesium chloride in the storage buffer. Finally, the extent of successful GTP loading into the GTPase has not consistently been assessed (**Table 1**). We also realized that a systematic analysis of conditions to achieve optimal GTP loading had not been performed.

**Table 1.**
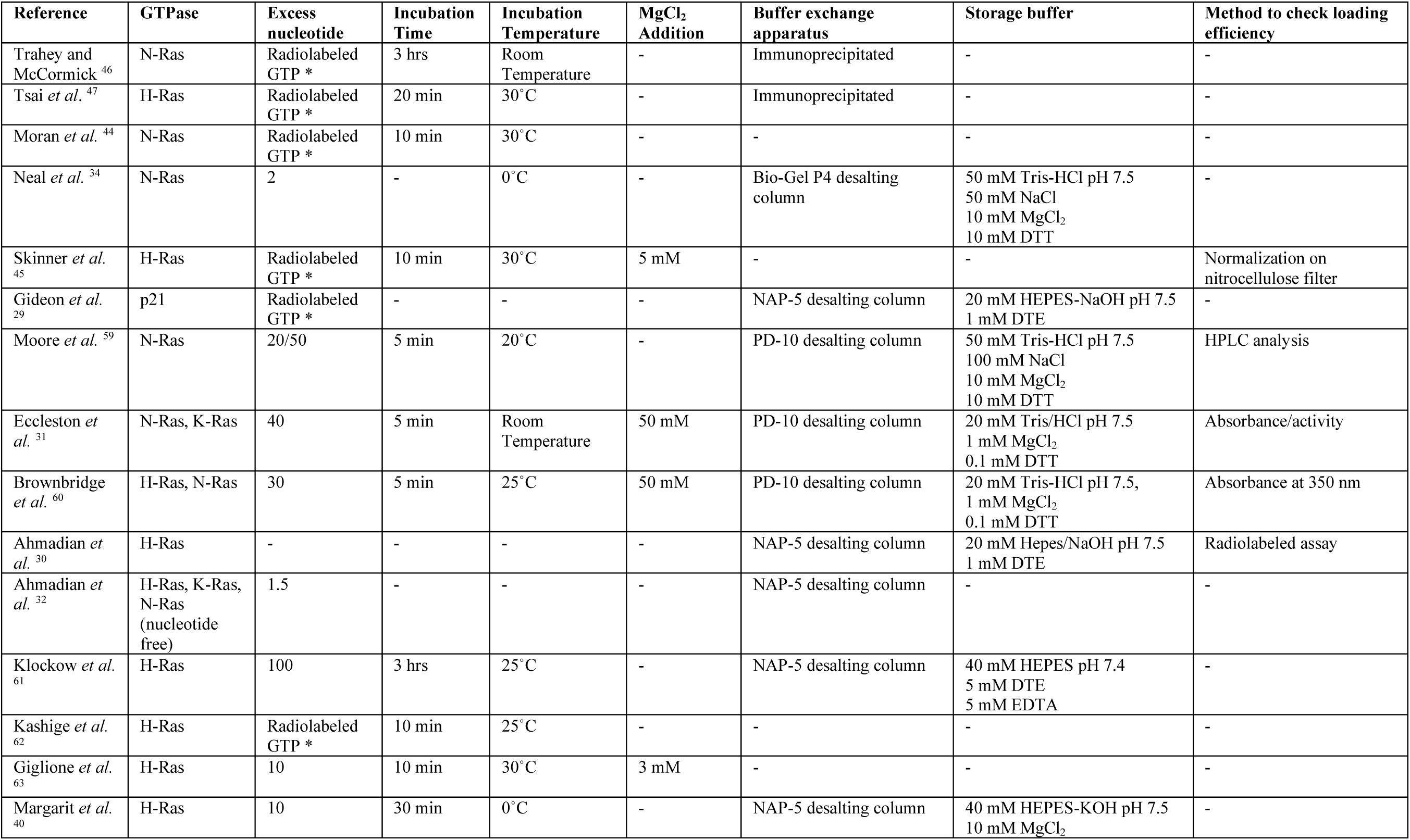
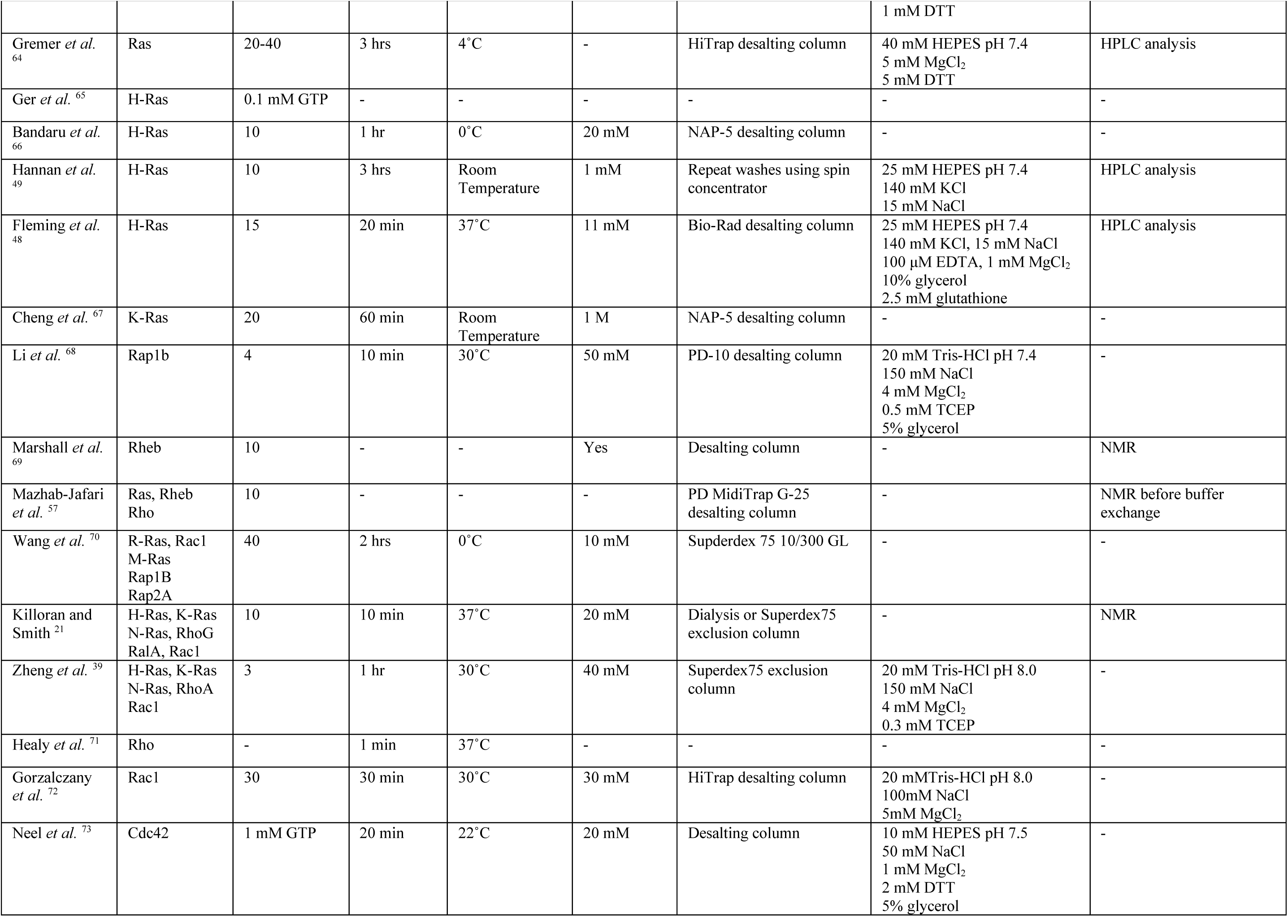
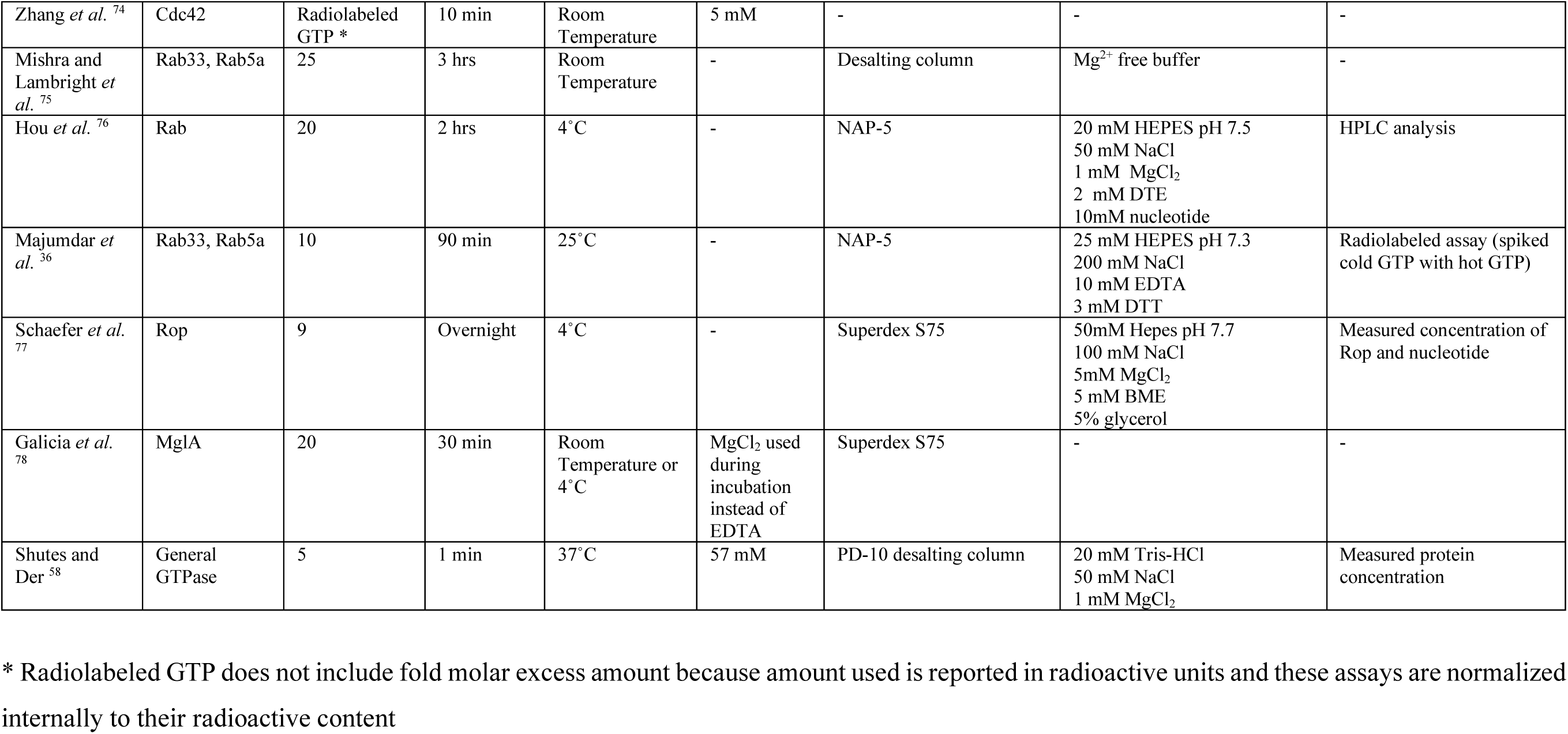
Experimental conditions for GTPase loading with GTP. Conditions used to load GTPases and assess nucleotide loading efficiency in previous studies. Excess nucleotide indicates fold excess over protein concentration. DTE indicates dithioerythritol and DTT indicates dithiothreitol. BME indicates β-Mercaptoethanol. EDTA indicates Ethylenediaminetetraacetic Acid.

We decided to conduct nucleotide exchange using conditions that we considered standard among published protocols (**Table 1**, **Figure 2A**). We incubated the purified G-domain of H-Ras in a solution containing 20 mM Tris pH 8, 100 mM NaCl, 10 mM EDTA, and 10-fold molar excess GTP. Following incubation for one hour at room temperature, we performed size exclusion chromatography to adequately separate unbound nucleotide from nucleotide-bound Ras (**Figure 2B**). To assess loading efficiency, we added EDTA, boiled, and sedimented denatured Ras leaving unbound nucleotide in the supernatant. Heating to 95°C on this timescale does not lead to intrinsic nucleotide hydrolysis. We then conducted anion exchange chromatography and identified GDP and GTP peaks according to separate evaluations of our nucleotide standards (**Figure 2C**). During loading, occasionally a GMP peak is observed, in accordance with previous literature, which we confirm with a GMP standard when necessary ^48, 49^ (**Supplemental Table 1**). We consistently find that using the above conditions to exchange GTP for GDP yield fewer than two thirds (62±3 %) of the Ras molecules loaded with GTP. This surprised us and we wondered whether it might be possible to optimize the amount of GTP bound to Ras.

**Figure 2.**
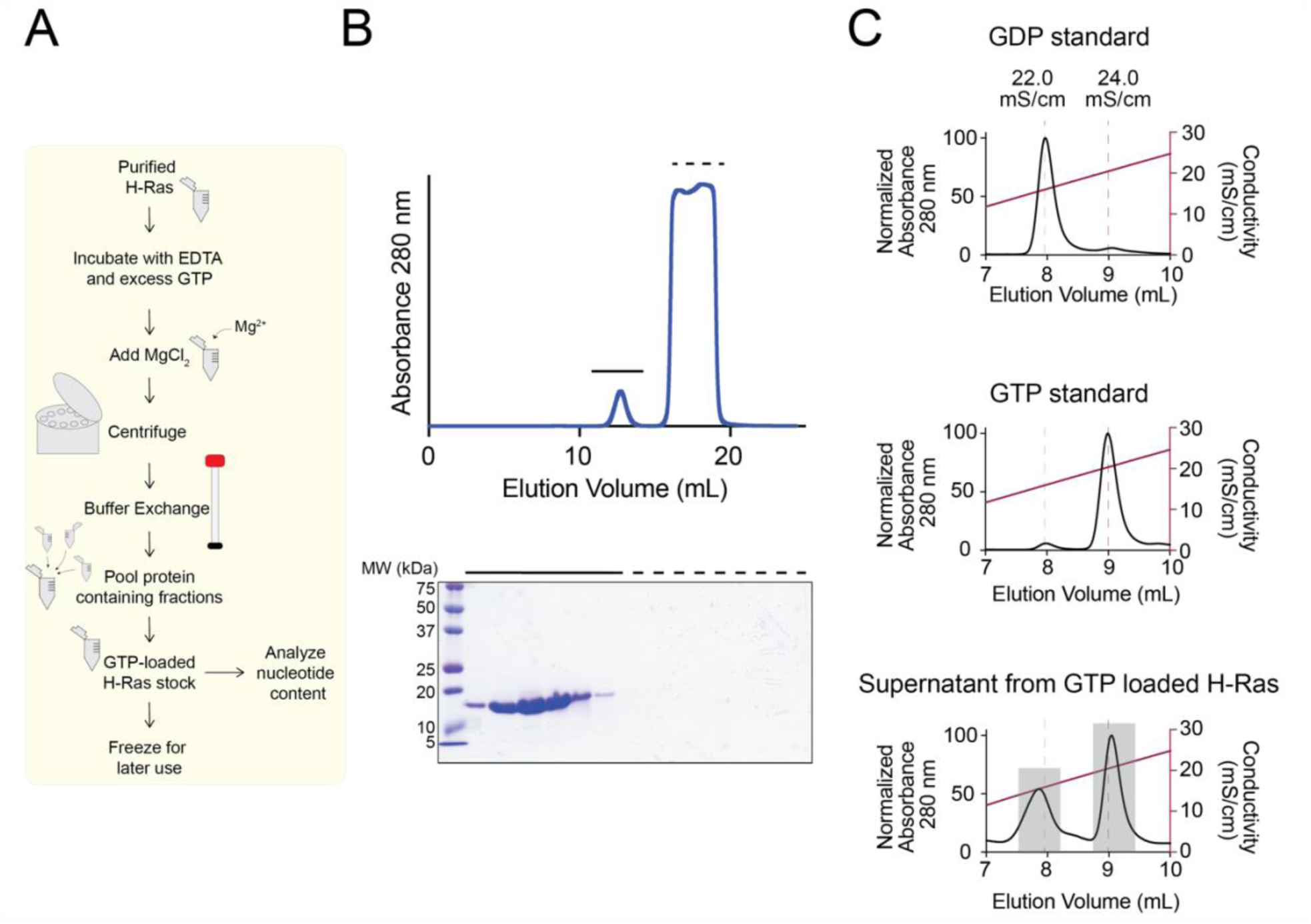
H-Ras loading using standard conditions. **A)** Flow chart of H-Ras loading. Purified H-Ras is incubated with EDTA and excess GTP. At the end of the incubation, Mg^2+^ is added in the form of MgCl_2_ to encourage nucleotide binding. Following centrifugation, buffer exchange using size exclusion chromatography is conducted to remove excess unbound nucleotide. H-Ras fractions are pooled and frozen for later use. **B)** Example buffer exchange during GTP loading. After incubating with EDTA and excess GTP, H-Ras was buffer exchanged to remove excess nucleotide by size exclusion chromatography using a Superdex 75 10/300 GL column in 20 mM Tris pH 8, 100 mM NaCl, 10 mM EDTA. The first peak is H-Ras protein and the second peak is unbound nucleotide as shown by the 15% SDS-PAGE. **C)** Nucleotide content of H-Ras after loading with GTP. After loading with GTP by incubating in EDTA and a 10-fold excess of GTP for one hour at room temperature H-Ras is only 55% GTP bound. Gray boxes indicate the peaks from which area under the curve was calculated.

We began by assessing whether temperature, time, or concentration of Ras might provide an improvement in GTP loading. In our analysis of previous studies, we observe that experimental loading time has been varied by orders of magnitude, with the shortest incubation at one minute and the longest at overnight (**Table 1**). We therefore assessed the GTP content of Ras when incubated with EDTA and nucleotide for 10, 30, and 60 minutes at 37°C and we observe a percent GTP loading ranging from 58±3 % to 66±1 %. While there is a statistical significance observed between 10 and 30 minutes, this is not maintained between 10 and 60 minutes, therefore we conclude time does not have an impact on loading efficiency (**Figure 3A, Supplemental Table 1**). Similarly, temperature is a factor that has been widely varied, with incubations performed at temperatures between 0°C (on ice) and 37°C (**Table 1**). We wondered if similar variations might alter loading of Ras with GTP and so measured the GTP content of Ras incubated for one hour on ice, room temperature, or 37°C. Similar to incubation time, temperature has no effect on Ras-GTP content with percent GTP loading ranging from 63±3 % to 68±4 %, (**Figure 3B, Supplemental Table 1**). Finally, we assessed whether altered concentration of Ras might impact loading efficiency. We compared the loading efficiencies for incubations where Ras was held at concentrations of 0.025 mM or 0.26 mM. Using a 10-fold excess of GTP concentration (0.25 mM or 2.6 mM GTP, respectively) we find no significant change in loading efficiency when varying the GTPase concentration, with percent GTP loading for Ras at 0.025 mM measured at 66±5 % and for Ras at 0.26 mM measured at 68±8 % (**Figure 3C, Supplemental Table 1**).

**Figure 3.**
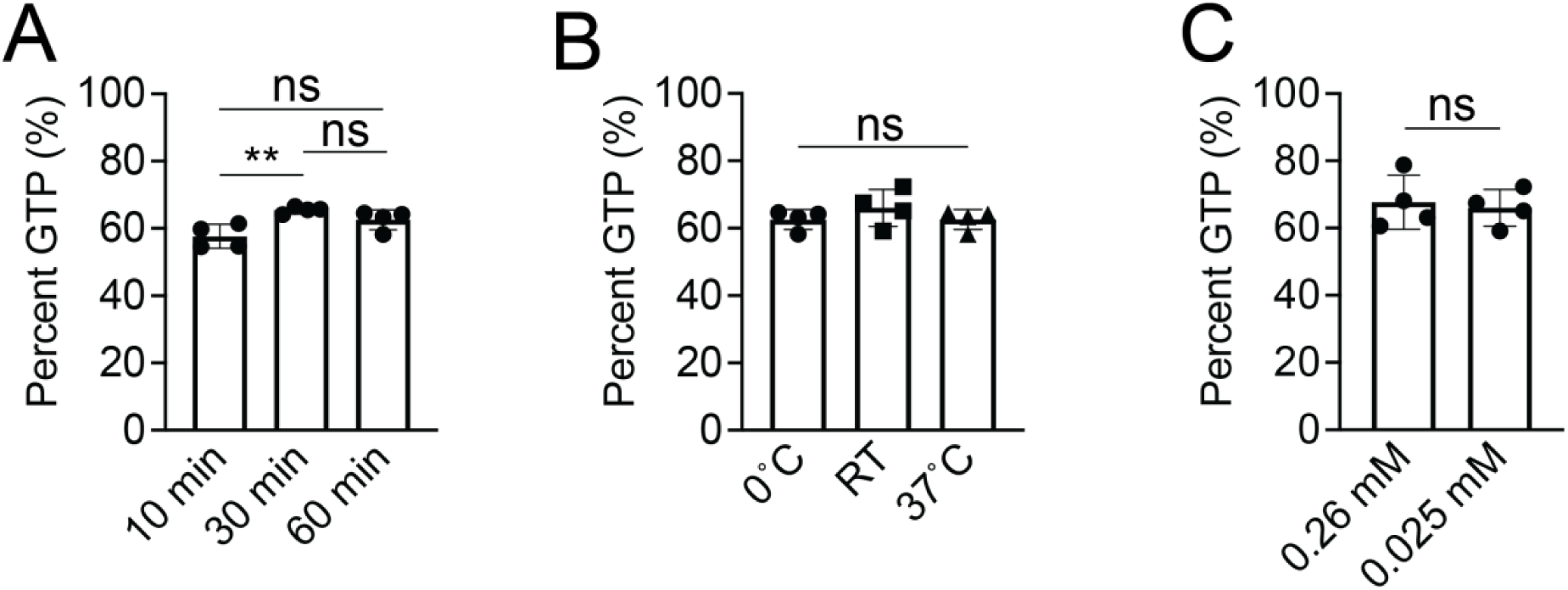
Time, temperature and H-Ras concentration do not impact GTP loading efficiency. **A)** Effect of time on H-Ras loading. 0.025 mM H-Ras was incubated with EDTA for 10, 30, or 60 minutes at 37°C before buffer exchange using a Superdex S75 10/300 column (Cytiva). Trials 1-12 from Supplemental Table 1 were used for these comparisons. n = 4. Error bars represent S.D. Statistical significance tested by one-way ANOVA p = 0.0083, F = 8.544, 2 degrees of freedom. Tukey’s multiple comparisons tests performed between the first column and subsequent columns. ** indicates p = 0.0069. ns indicates not significant. **B)** Effect of temperature on loading efficiency. 0.025 mM H-Ras was incubated with EDTA for one hour at either 0°C (on ice), room temperature, or 37°C before buffer exchange using a Superdex S75 10/300 column (cytiva). Trials 9-20 from Supplemental Table 1 were used for these comparisons. Loading efficiency was analyzed using anion exchange chromatography. n = 4. Error bars represent S.D. Statistical significance tested by one-way ANOVA p = 0.4047, F = 1.002, 2 degrees of freedom. Tukey’s multiple comparisons tests performed between the first column and subsequent columns. ns indicates not significant. **C)** Effect of concentration of H-Ras on loading efficiency. Either 0.025 mM or 0.26 mM H-Ras were loaded with respective 10-fold excess GTP and loading efficiency analyzed using anion exchange chromatography. Trials 13-16 and 21-24 from Supplemental Table 1 were used for this comparison. n = 4. Error bars represent S.D. An unpaired t test was performed and p = 0.7419. ns indicates not significant. Percent GTP was calculated for all trials by integrating the area under the curve for each (GTP or GDP) nucleotide peak.

Next, we assessed the role of magnesium on GTP loading. Small GTPases use magnesium ions to coordinate the phosphate groups of the bound nucleotide, therefore addition of MgCl_2_ is sometimes used to quench EDTA at the end of the incubation, restoring an environment more favorable to nucleotide-binding ^50–52^. Nonetheless, in our analysis of previous studies we find that magnesium addition is not consistently performed (**Table 1**) potentially because of concern that magnesium addition may help initiate GTP hydrolysis ^16, 53^. We therefore asked whether magnesium addition aids nucleotide exchange by comparison of GTP content for samples that were or were not treated with addition of MgCl_2_ at the conclusion of incubation with EDTA. We find GTP loading for samples incubated with 10 mM EDTA to be 64±7 %, but for samples quenched with 15 mM MgCl_2_ at the end of the incubation with EDTA the percent of GTP loading rose significantly to 75±9 % (**Figure 4A, Supplemental Table 1**).

**Figure 4.**
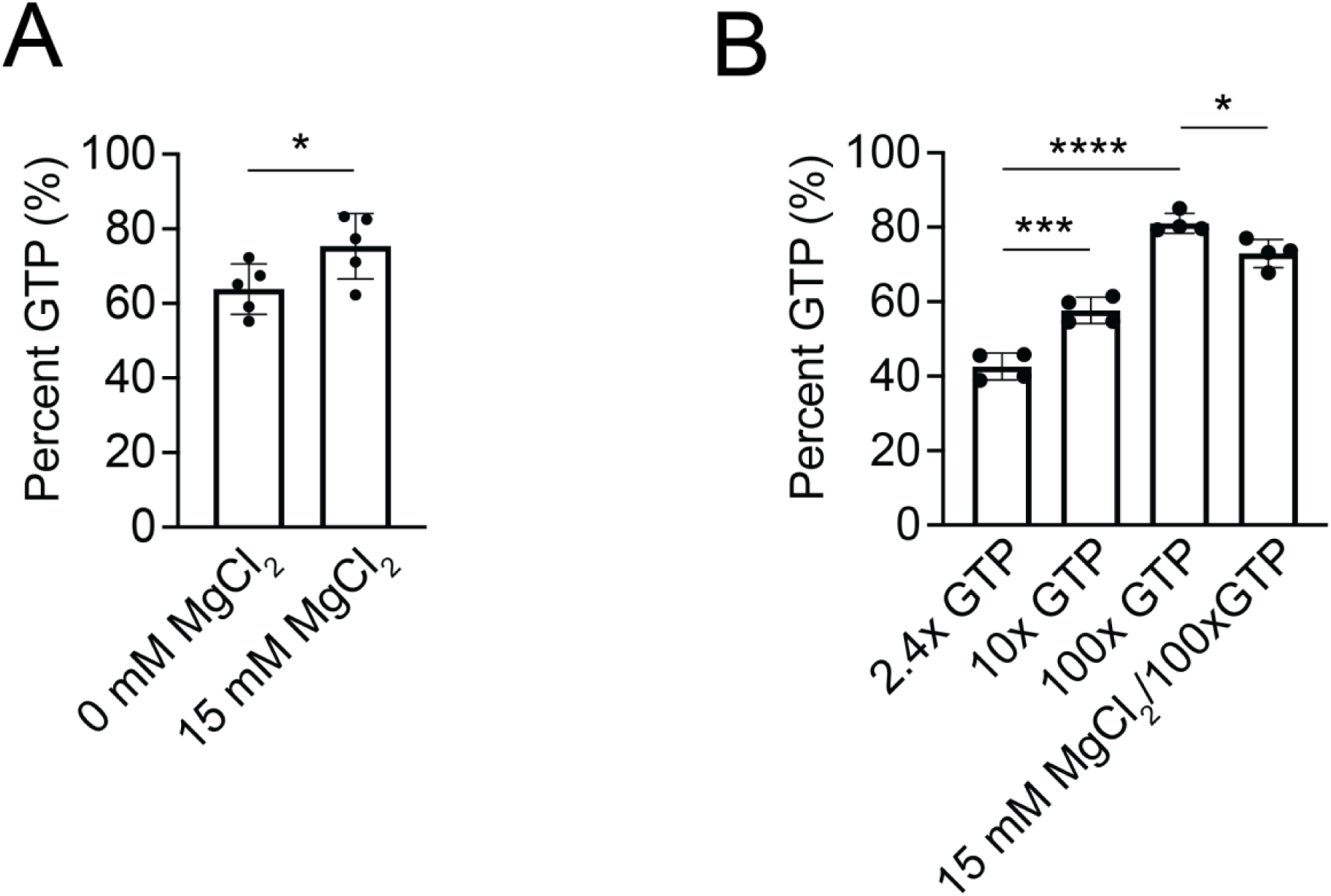
Increased excess GTP improves GTP loading efficiency. **A)** Effect of magnesium addition on GTP loading efficiency. H-Ras was loaded with 10-fold excess GTP in the presence of EDTA for 1 hour at room temperature. At the end of the ETDA incubation period, 15 mM MgCl_2_. Loading efficiency was analyzed using anion exchange chromatography. Trials 13-16 and 26-31 from Supplemental Table 1 were used in this comparison. n = 5. Error bars represent S.D. Statistical significance was tested using an unpaired t test and p = 0.0488. **B)** Effect of GTP excess on GTP loading efficiency. 100-fold molar excess GTP leads to better loading. 0.025 mM H-Ras was loaded using either 2.4-fold, 10-fold, or 100-fold molar excess GTP and loading efficiency was analyzed using anion exchange chromatography. Combining 100x molar excess with an addition of magnesium at the end of the EDTA incubation did not significantly alter loading efficiency. Trials 1-4 and 32-43 from Supplemental Table 1 were used for this comparison. n = 4. Error bars represent S.D. Statistical significance tested by one-way ANOVA p = <0.0001, F = 98.06, 3 degrees of freedom. Tukey’s multiple comparisons tests performed between the first column and subsequent columns. *** indicates p = 0.0002. **** indicates p = <0.0001. * indicates p = 0.0260. Percent GTP was calculated for all samples by integrating the area under the curve for each (GTP or GDP) nucleotide peak.

Analysis of previous work also suggests variations in the amount of GTP used for incubation. Some previous studies used GTP in a molar excess of only 1.5-fold, but others use molar excess up to 100-fold. Typically, however, studies have used GTP molar excess in the range of 10-fold over protein concentration. As these concentrations are all many fold higher than the K_d_ of GTP for Ras ^18, 22^, we assumed no impact would be observed on percentage of GTP loaded. Nonetheless, we compared the impact of 2.4-fold, 10-fold and 100-fold excess GTP concentration over the concentration of Ras. Surprisingly, we find that increasing the amount of excess GTP present during incubation can significantly increase loading efficiency. For our experiments using 2.4-fold excess we observe GTP loading of 42±4 %, using 10-fold excess we observe GTP loading of 58±4 %, and using 100-fold excess we observe GTP loading for 80±1 % of Ras (**Figure 4B, Supplemental Table 1**). Large excesses of GTP nucleotide significantly facilitates GTP loading.

Finally, to consolidate our observed increases in GTP loading, we assessed the effect of both MgCl_2_ addition at the conclusion of incubation and incubation with GTP at 100-fold excess to the concentration of Ras. We observe that GTP is loaded into 72±4 % of Ras molecules (**Figure 4B, Supplemental Table 1**), a significant decrease in GTP loading compared to without MgCl_2_. We interpret these results to conclude that the determining factor of loading efficiency is fold excess of GTP nucleotide concentration over GTPase, and that addition of MgCl_2_ does not significantly impact GTP loading, although the addition of MgCl_2_ at lower fold excess amounts of GTP may be beneficial for GTP loading.

### An EDTA-facilitated loading technique is applicable to all Ras isoforms

We next wondered if this loading protocol would be applicable to the other Ras isoforms since they all have nearly identical active sites (**Figure 5A, Supplemental Table 1 and 2**). To test if our loading protocol is applicable to the other Ras isoforms, we purified N-Ras and K-Ras from bacteria using the same protocol and compared the loading efficiencies of H-, K-, and N-Ras under the same conditions (10-fold excess GTP, 10 minutes, 37°C). All the isoforms had approximately the same loading efficiency, with only a slight difference observed between K-Ras and N-Ras. Overall, this implies our protocol is applicable to all Ras isoforms, consistent with the observation all isoforms contain the same conserved residues in their active sites (**Figure 5B, Supplemental Table 1 and 2**).

**Figure 5.**
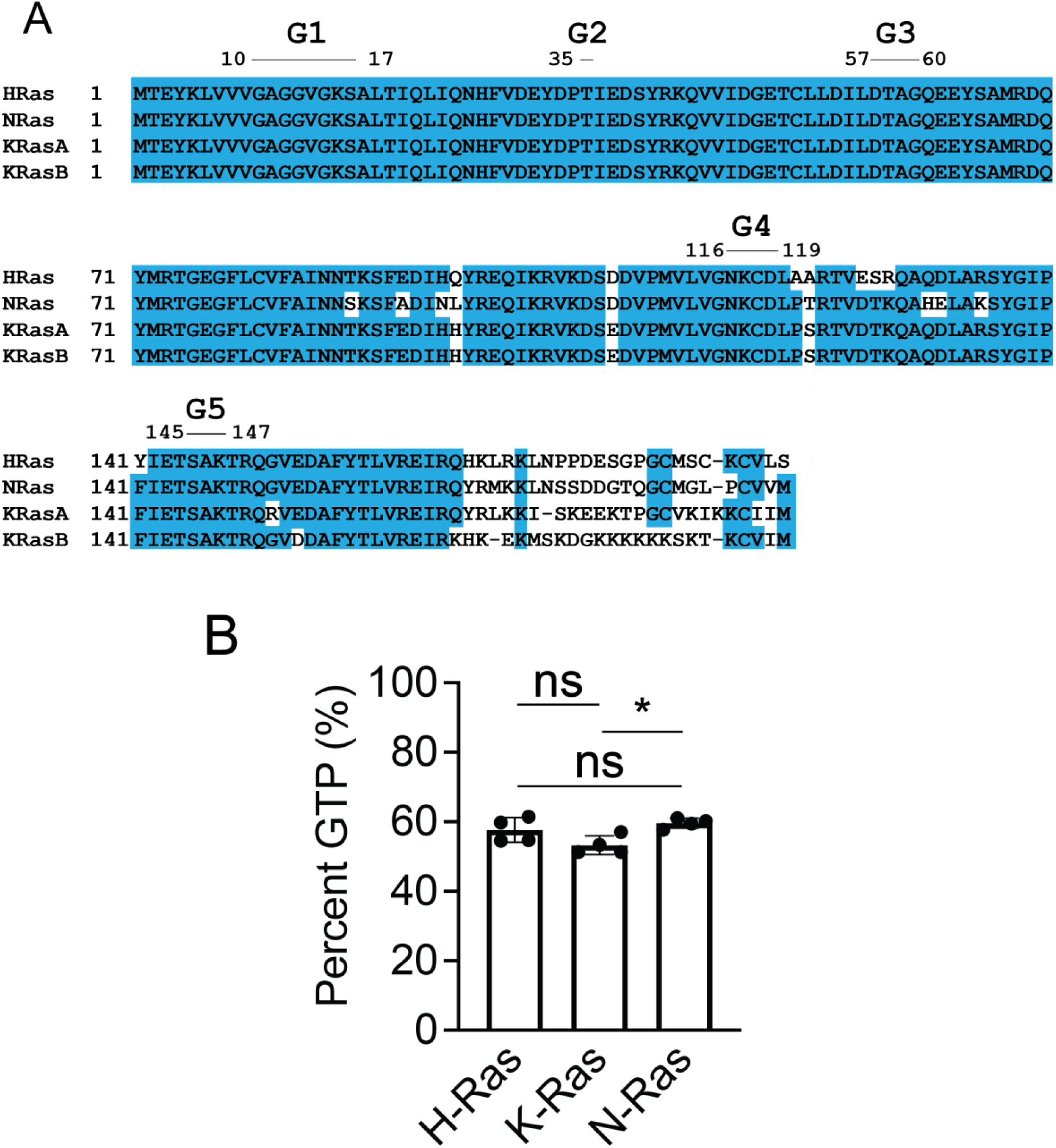
Loading with GTP is consistent across Ras isoforms. **A)** Sequence alignment of Ras isoforms. (H-Ras Uniprot ID P01112; N-Ras Uniprot ID P01111; K-Ras Uniprot ID P01116) Blue represents sequence conservation. The functional G motifs necessary for GTP binding are indicated. MAFFT online was used to align the isoforms.^79^ **B)** Comparison of GTP loading for Ras isoforms. 0.025 mM Ras was loaded using 10x GTP for 10 minutes at 37°C. Trials 1-4 from Supplemental Table 1 and 98-105 from Supplemental Table 2 were used in this comparison. n = 4. Error bars represent S.D. Statistical significance tested by one-way ANOVA p = 0.0237, F = 5.835, 2 degrees of freedom. Tukey’s multiple comparisons tests performed between the first column and subsequent columns. * indicates p = 0.0215. ns indicates not significant.

### Storage buffer for GTP-loaded Ras should contain EDTA but not magnesium

After nucleotide exchange it is common to remove excess nucleotide. In our survey of previous studies we find that this is usually accomplished by a buffer exchange step via a desalting column or size exclusion chromatography, but that the contents of the destination buffer (which we term the ‘storage buffer’) shows some variability (**Table 1**). Importantly, there is discrepancy between whether to add EDTA or magnesium to the storage buffer. We therefore assessed the impact of inclusion of either 10 mM EDTA, or 10 mM MgCl_2_, or both 10 mM EDTA and 10 mM MgCl_2_, in a storage buffer of 20 mM Tris pH 8, 150 mM NaCl when we conduct size exclusion chromatography buffer exchange at 4 °C. We find that addition of EDTA to the storage buffer has no effect on the amount of GTP loaded compared to buffer alone (63±1 % and 63±3 % for buffer alone and 10 mM EDTA, respectively) (**Figure 6A, Supplemental Table 1**). In contrast, when we include MgCl_2_ in the storage buffer, we find a significant decrease in bound GTP (50±9 %) occurs on the timescale of buffer exchange (**Figure 6A, Supplemental Table 1**). Addition of both EDTA and MgCl_2_ in the storage buffer results in further decrease of bound GTP (36±1 %), potentially due to increased cycling through hydrolysis, nucleotide release, and nucleotide binding facilitated by the presence of EDTA in addition to MgCl_2_. This suggested to us that although addition of MgCl_2_ and excess GTP provide optimal conditions for achieving nucleotide exchange, extended exposure to MgCl_2_ may facilitate activity of the enzyme and accelerate cleavage of GTP into GDP and inorganic phosphate.

**Figure 6.**
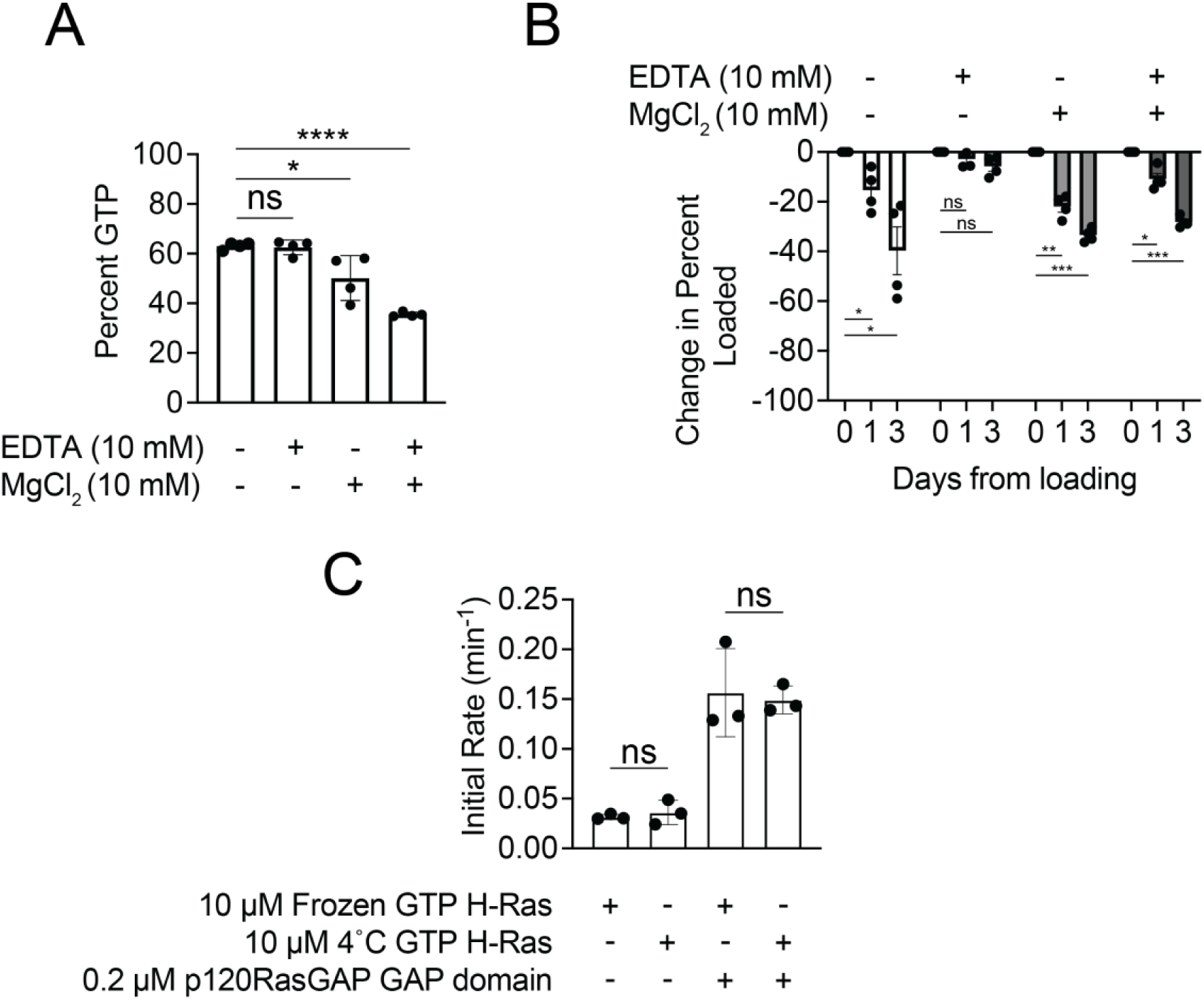
Effect of storage buffer on loading efficiency. **A)** Effect of storage buffer on loading efficiency following buffer exchange. H-Ras was loaded with 10-fold excess GTP in the presence of EDTA. After the incubation period, H-Ras was buffer exchanged into 20 mM Tris pH 8, 150 mM NaCl, or this buffer containing 10 mM MgCl_2_ and/or 10 mM EDTA. Trials 17-20 and 44-55 from Supplemental Table 1 were used in this comparison. n = 4. Statistical significance tested by one-way ANOVA p = <0.0001, F = 28.94, 3 degrees of freedom. Tukey’s multiple comparisons tests performed between the first column and subsequent columns. * indicates p = 0.0121. **** indicates p = <0.0001. ns indicates not significant. **B)** Long-term effect of storage buffer at 4°C on loading efficiency. After loading H-Ras with GTP, H-Ras was buffer exchanged into 20 mM Tris pH 8, 150 mM NaCl buffer either including or excluding 10 mM EDTA and/or MgCl_2_ as indicated. The sample was stored at 4°C and the nucleotide content was analyzed over time. All samples are normalized according to initial GTP content, and the amount decrease of GTP from the day of original loading was plotted. Trials 1-3, 32-33, 44, and 56-97 from Supplemental Table 1 were used for this comparison. Loading efficiency was analyzed using anion exchange chromatography. n = 4. Statistical significance tested by two-way ANOVA. Row factor p = 0.0002, F = 104.1, 2 degrees of freedom. Uncorrected Fisher’s LSD tests performed between different days per condition tested. * indicates p = 0.0348, 0.0259, and 0.0164 as they appear left to right in figure. ** indicates p = 0.0022. *** indicates p = 0.0001. ns indicates not significant. **C)** Initial rates of GTP H-Ras either frozen immediately after GTP loading or incubated at 4°C for 3 days. GTP H-Ras was buffer exchanged into storage buffer containing 20 mM Tris pH 8, 150 mM NaCl, 10 mM EDTA and either frozen or stored at 4°C for three days. Then GTPases were performed using 10 µM GTP H-Ras, 15 µM Phosphate Sensor and 0.2 µM GAP domain when indicated. n = 3 representing the average of two technical duplicates. Error bars represent S.D. Statistical significance tested by one-way ANOVA p = 0.0002, F = 24.51, 3 degrees of freedom. Tukey’s multiple comparisons tests performed between the first column and subsequent columns. ns indicates not significant.

We therefore considered that excess MgCl_2_ may also prove deleterious for long-term storage of GTP-loaded enzyme. To assess this, we compared the long-term effect of storing GTP-loaded H-Ras in a buffer of 20 mM Tris pH 8, 150 mM NaCl with storage buffer containing 10 mM EDTA, 10 mM MgCl_2_, neither, or both. Following buffer exchange, we assessed the amount of H-Ras bound after one and three days of storage at 4°C. We observe that over a period of three days storage without the addition of EDTA is detrimental to maintenance of bound GTP (**Figure 6B)**. In our experiments, the only samples with minimal nucleotide hydrolysis were those stored in 10 mM EDTA and no magnesium. To ensure this is due to reduced hydrolysis activity and not destabilization of the H-Ras protein we conducted GTPase assays using GTP H-Ras stored frozen or at 4°C immediately after loading for three days. We observe there is no difference in initial rate of phosphate release between the frozen and 4°C incubated H-Ras for either intrinsic H-Ras phosphate release and GAP assisted H-Ras phosphate release (**Figure 6C**). Our results indicate that optimal long-term storage of GTP-loaded H-Ras to limit enzymatic activity can be achieved by buffer exchange into storage buffer with EDTA but without magnesium.

### Impact of cancer mutations on nucleotide bound upon purification

Alterations in Ras genes are associated with significant populations of cancers, with residues Gly12, Gly13 and Gln61 representing the most frequent sites of mutation. We wondered whether our protocol might be useful for assessing the extent of bound GTP in recombinantly expressed H-Ras G-domain when each of these mutations are present. According to previous literature, for H-Ras the most commonly observed mutations at codons 12 and 13 are G12V and G13R, and at codon 61 the most well studied mutation is Q61L ^9, 10, 12, 13, 54, 55^. Therefore, we chose to assess these mutations. We observe that all three mutant H-Ras proteins co-purify with significantly more GTP bound that wild-type Ras, with wild-type G12V and Q61L containing 29±6 % and 32±2 % GTP, and G13R with 12±2 % GTP (**Figure 7A, Supplemental Table 3**). This is likely due to the decreased intrinsic GTP hydrolysis rate observed with the mutant H-Ras molecules ^21, 25^. We noted that these samples were purified but not subjected to EDTA incubation, consequently a portion is bound to GMP, a feature that has previously been observed ^48, 49^. We then assessed whether we could exchange bound GDP for GTP in the context of the G12V mutation. We decided to load with 10-fold excess GTP to allow for a wider dynamic range. Interestingly, we observe that under the same conditions G12V mutant H-Ras is significantly less efficient at loading GTP than wild type H-Ras with GTP loading of 36±9 % for the G12V mutant and 58±4 % for wild type H-Ras. Therefore, EDTA loading seems to have only a minor impact on the amount of bound GTP (**Figure 7BC, Supplemental Table 4)**.

**Figure 7.**
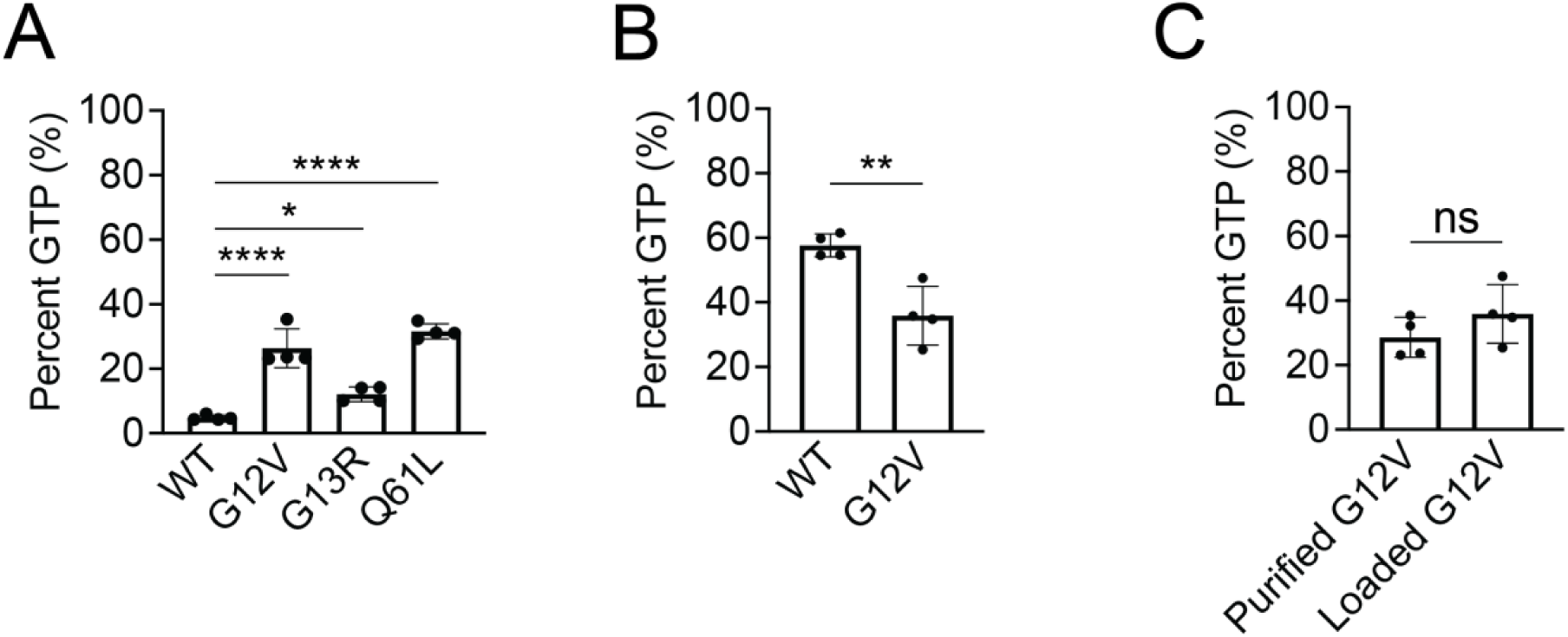
Assessment of nucleotide composition for recombinantly expressed H-Ras cancer mutants. **A)** Nucleotide content of recombinantly expressed H-Ras cancer mutations. Analysis of purified samples from Supplemental Table 3 were used in this comparison. Error bars represent S.D., n = 4. Statistical significance tested by one-way ANOVA p = <0.0001, F = 51.68, 3 degrees of freedom. Tukey’s multiple comparisons tests performed between the first column and subsequent columns. * indicates p = 0.0482. **** indicates p = <0.0001. **B)** Comparison of GTP loading for wild type and G12V H-Ras. 0.025 mM H-Ras-G12V was loaded using 10x GTP for 10 minutes at 37°C and compared to wild type H-Ras loaded under the same conditions. Trials 1-4 from Supplemental Table 1 and 106-109 from Supplemental Table 4 were used in this comparison. n = 4. Error bars represent S.D. Statistical significance tested via an unpaired t test p = 0.0042. **C)** Comparison of the GTP content of purified and GTP-loaded H-Ras-G12V. Purified samples from Supplemental Table 3 and trials 106-109 from Supplemental Table 4 were used for this comparison. n = 4. Error bars represent S.D. Statistical significance tested via an unpaired t test p = 0.2312. Percent GTP was calculated by integrating the area under the curve for each (GTP or GDP) nucleotide peak.

## Discussion

In this study we assessed the previous literature for consensus methods to conduct H-Ras *in vitro* nucleotide exchange, yet observed some variance. We therefore conducted a careful biochemical analysis of conditions that achieve maximal exchange of nucleotide in purified Ras and used anion exchange chromatography to monitor the bound nucleotide composition. We continued by demonstrating optimized conditions for storage of GTP loaded H-Ras. To see if this technique is applicable to all Ras isoforms, we compared the loading efficiency of H-Ras, K-Ras, and N-Ras under the same conditions and saw similar loading efficiencies. We concluded our study with a demonstration of the utility of this method for assessment of the nucleotide binding state of recombinantly produced H-Ras containing common cancer-associated mutations. We therefore provide new insights into best practices for *in vitro* biochemical analysis of Ras proteins.

Key for this study was the use of anion exchange chromatography to assess nucleotide composition of the Ras samples. In prior literature for small GTPase *in vitro* nucleotide exchange composition analysis has sometimes been achieved by reverse-phase chromatography ^49^, but there has been no consensus formed for a standard and easily accessible method (**Table 1**). Here, we used anion exchange chromatography as a tool for determination of nucleotide content of purified and loaded Ras and build on the previous work of our, and other, groups ^41–43^. By denaturing the protein, we found that nucleotide is released into solution and can be monitored by supernatant analysis over anion exchange chromatography by standard equipment. Importantly, because guanine nucleotides absorb significantly at both 254 nm and 280 nm wavelengths, we demonstrated quantitative assessment of the nucleotide composition (GDP versus GTP) by integration under elution curves from anion exchange chromatography at either 254 nm or 280 nm. These wavelengths are accessible to most common FPLC systems, so we believe that use of standard FPLC equipment to monitor nucleotide composition of purified small GTPases can become a commonly performed technique for Ras, other small GTPases, and potentially other nucleotide-binding proteins.

EDTA treatment of GDP-loaded small GTPases to facilitate nucleotide exchange has extensive precedence ^34, 44–47^ (**Table 1**). Nonetheless, there are variations in reported experimental protocols to achieve optimal nucleotide exchange, and our analysis of the prior literature led us to assess an array of variables that have been frequently altered. We found that ranges used for multiple of the reported variables do not impact GTP loading for Ras, these include incubation time, temperature, and concentration of H-Ras in the experiment. In contrast, we found that although there was no consensus in previous literature on whether or not to include addition of magnesium at the conclusion of EDTA incubation, in our hands magnesium addition shows only qualitative changes in GTP-loading of H-Ras. Similarly, the amount of excess GTP present has not been consistent, but we find that yield was significantly improved by increasing amounts of excess GTP present. Our study therefore provides guidance for experimental protocols for nucleotide exchange.

Experiments subsequent to nucleotide exchange often require excess nucleotide to be removed, achieved by buffer exchange. For this, our literature search revealed experimental variability in the inclusion of magnesium in the destination ‘storage buffer’. Experimentally, we found that magnesium in the storage buffer was deleterious to GTP loading of Ras, and that when stored at 4°C the presence of magnesium results in extensive loss of GTP-bound GTP over just three days. Instead, our analyses indicate that optimal conditions for storage of GTP-loaded Ras are in the presence of EDTA and absence of magnesium. We conclude that care should be taken to avoid storage of GTP-loaded small GTPases in the presence of magnesium, and if storage at 4°C is required an FPLC analysis of GDP/GTP composition should be performed prior to experimentation.

Finally, we used this method to test the nucleotide-bound state of recombinantly expressed H-Ras containing common cancer-associated mutations. The mutant H-Ras proteins are purified in the same way as wild type by a four-day purification scheme. We show that the cancer mutants purify from *E. coli* with significantly more GTP content than wild type H-Ras and demonstrate a distinction between the mutants where we observe that G13R purifies from *E. coli* with less bound GTP than either G12V or Q61L. This agrees with previous work showing that Ras cancer mutants purify with more GTP bound, and that the intrinsic GTP hydrolysis activity of K-Ras is decreased in all three mutants (G12, G13, Q61), but less of an impact on GTP hydrolysis is observed for G13 compared to mutations at G12 and Q61^49, 56^. We note, however, that none of the cancer-associated mutations of H-Ras purify from recombinant expression in *E. coli* as 100% GTP-bound states, in accordance with previous work ^49^. We also demonstrated that the G12V mutant of H-Ras was resistant to nucleotide exchange. This may hint that other GTPases are differently amenable to this technique, and may be evidenced by one group adding urea to the exchange buffer when trying to load Rho with GTP^57^.

In this study we show the use of EDTA-facilitated method GTP loading of H-Ras can be optimized to achieve loading of approximately 80%. Nonetheless, an array of previous literature claims complete GTPase loading with similar methodology (e.g. **Table 1**). We posit that complete loading may not have been achieved in some of these studies, but we also provide an accessible solution – the use of FPLC analysis to assess GTP loading. Overall, our study provides experimental insights into optimal conditions for the use of EDTA-facilitated nucleotide exchange for small GTPases, demonstration of the use of anion exchange chromatography to quantitatively assess nucleotide composition of Ras, and an assessment of the impact of cancer mutations on nucleotide composition.

## Methods

### Purification of H-Ras and GAP constructs

All Ras proteins were purified as described in ^37^. Briefly, the sequence containing residues 1-167 of human H-Ras (UniProt ID: P01112) were cloned into a modified pET vector containing a 6xHis-tag and TEV cut site using restriction enzymes BamHI and XhoI. Cancer mutations were introduced to the plasmid by QuikChange (Agilent). N-Ras (#25256) and K-RasB (#25153) isoforms were gifts from Cheryl Arrowsmith (Addgene plasmid # 25256 ; http://n2t.net/addgene:25256 ; RRID:Addgene_25256; and Addgene plasmid # 25153 ; http://n2t.net/addgene:25153 ; RRID:Addgene_25153). These plasmids contain N-Ras residues 1-172 and K-RasB 1-169 in pET His vectors with a TEV cut site. The Q61H mutation in the K-RasB clone was reverted back to wild type using Agilent’s Quikchange kit. Purification of these constructs was identical to purification of H-Ras constructs. The plasmid was transformed into Rosetta (DE3) cells and the cells subsequently grown at 37°C shaking to an O.D. of 0.6-0.8 when protein production was induced via addition of 0.25 mM Isopropyl β-d-1-thiogalactopyranoside (IPTG). Once induced, the culture was transferred to 18°C and incubated overnight shaking. Cells were harvested by centrifugation at 2000 xg for 30 minutes and resuspended in lysis buffer containing 50 mM HEPES pH 8, 500 mM NaCl. Cells were lysed by sonication after addition of 25 μg/ml lysozyme and multiple freeze-thaw cycles. The lysate was clarified by spinning at 5000 xg for 1 hour at 4°C. Clarified lysate was then added to Ni-NTA agarose beads (Thermo Fisher Scientific) and rocked at 4°C for 1 hour to promote 6xHis-tag binding. H-Ras was eluted using a step wise gradient with 20 mM, 40 mM, 100 mM, 250 mM, and 500 mM imidazole in lysis buffer. Fractions containing H-Ras were then pooled and the 6xHis-tag removed by incubation with his-tagged tobacco etch virus protease (His-TEV). Imidazole concentration was reduced by dialyzing overnight against lysis buffer. To remove free tag and His-TEV, the protein was next added to Ni-NTA agarose beads for 1 hour, and the supernatant collected and concentrated. During concentration, 20 mM Tris pH 8.0 buffer was added to reduce the NaCl in preparation for anion exchange chromatography. Anion exchange chromatography was performed using a ResourceQ (1 ml, Cytiva) column with buffers A (20 mM Tris pH 8.0) and B (20 mM Tris pH 8.0, 1 M NaCl). Finally, fractions containing Ras were pooled and concentrated and subjected to size exclusion chromatography using a HiLoad 16/600 Superdex 75 prep grade (Cytiva) column and 20 mM Tris pH 8.0, 150 mM NaCl buffer. All Ras proteins were purified using the same method. All Ras proteins take four days to purify.

Residues corresponding to p120RasGAP’s GAP domain (714-1047) were also cloned into a modified pET vector containing a 6xHis-tag and TEV cut site using restriction enzymes BamHI and XhoI and purified using the same protocol as the Ras proteins.

### GTP loading and nucleotide analysis of H-Ras

Purified Ras (at concentrations 0.025 mM and 0.26 mM) was loaded by adding 10-100 fold excess GTP in buffer containing 20 mM Tris pH 8, 100 mM NaCl, and 10 mM ethylenediaminetetraacetic acid (EDTA). Ras was incubated at temperatures between 0°C (on ice) and 37°C for 10-60 minutes as indicated for each experiment. Relevant reactions were stopped by addition of 15 mM MgCl_2_ as indicated in the text. Reactions were then centrifuged at 16,900 xg at 4°C for 10 minutes before buffer exchange by application to a Superdex 75 10/300 GL Increase column in buffer indicated by the experiment (with all buffers containing 20 mM Tris pH 8.0 and 150 mM NaCl and sometimes containing 10 mM EDTA and/or 10 mM MgCl_2_). Loading efficiency was assessed by incubation of 0.01 nmol of protein sample with 10 mM EDTA at 95°C for 15 minutes. Denatured protein was pelleted by centrifugation at 16,900 xg at 4°C for 10 minutes. The supernatant containing the nucleotide was then diluted with 20 mM Tris pH 8 to reduce the NaCl concentration to less than 50 mM and applied to a MonoQ 5/50 GL (Cytiva) column to separate the nucleotide content. Peaks were compared to GDP and GTP standards and area under the curve was used to calculate percent GTP bound using the integration feature in Unicorn software (Cytiva) and integrated under the curve.

### Guanosine nucleotide standards

GMP, GDP and GTP standards were analyzed using the same method as the Ras samples on the monoQ 5/50 GL (Cytiva) column. Nucleotides were purchased from Millipore Sigma.

### GTPase assays

GTPase assays were performed using Phosphate Sensor purchased by Thermo Fisher Scientific to measure phosphate release. For these assays H-Ras was loaded in 10-50 fold excess GTP in the presence of EDTA. After incubating for 10 minutes at 37°C the reaction was stopped with the addition of 15 mM MgCl_2_. Excess nucleotide was removed with a buffer exchange step consisting of the application of the H-Ras to a Superdex 75 Increase column in 20 mM Tris pH 8, 150 mM NaCl, 10 mM EDTA. Half of each sample was frozen immediately after GTP loading and half was stored at 4°C for three days. After three days, the frozen sample was thawed and both thawed and 4°C incubated samples were spun down at 16,900 xg at 4°C for 10 minutes. The reaction was performed in buffer containing 20 mM Tris pH 8, 50 mM NaCl, 0.01% Triton-X, and 1 mM TCEP. Each reaction contained 15 µM Phosphate Sensor, 10 µM GTP H-Ras, and 0.2 µM GAP domain derived from p120RasGAP as indicated by the text. After three minutes of measuring the baseline fluorescence each reaction was started by the addition of 5 mM MgCl_2_. The reactions were read at the shortest minimum interval for 30 min Synergy H1 plate reader (BioTek Agilent) using Gen5 3.11 software (https://www.agilent.com/en/product/microplate-instrumentation/microplate-instrumentation-controlanalysis-software/imager-reader-control-analysis-software/biotek-gen5-software-for-detection-1623227). Data were then normalized to fluorescence as suggested by ^58^ and initial rates determined by measuring the slope during the first 10 min of assay time. Each *n* represents the average of two technical replicates.

### Statistical Assessment

Statistical significance was assessed by determining *p* values calculated by an unpaired t test when comparing two conditions, and a one-way ANOVA when comparing three or more conditions with Tukey’s multiple comparisons test, and a two-way ANOVA when comparing multiple days of multiple conditions. The test used for each comparison is indicated in the figure legends (GraphPad Prism v. 10.1.1) (https://www.graphpad.com/).

## Data Availability

Raw data for the chromatograms is available as an excel spreadsheet which can be obtained by request to the corresponding author (Titus J. Boggon).

## Author Contributions

Kimberly J. Vish: conceptualization, investigation, data curation, writing - original draft, writing - review and editing; Asha P. Rollins: investigation, writing – review and editing; Maxum E. Paul: investigation, writing - review and editing; Titus J. Boggon: conceptualization, funding acquisition, writing - review and editing.

## Declarations of Interest

The authors declare no competing interests.

## Acknowledgements

We thank Amy Stiegler Wyler, Moitrayee Bhattacharyya, and Benjamin Turk for helpful discussions. We thank Cheryl Arrowsmith for the gifts of KRASB and NRASA. K.J.V. supported by T32GM008283 and F31HL165968. M.E.P. supported by T32GM007324 and F31HL16758. This research was supported by R01NS117609 to T.J.B.

## Notes

### Competing Interest Statement

The authors have declared no competing interest.

