## Supplemental Information for "Optimized conditions for GTP loading of Ras"

**Supplemental Table 1. Conditions used for all loading replicates.** Depending on the condition being tested some replicates were used in multiple figures as indicated by the figure legends.

| Trial # | $Mg^{2+}$ Addition | Amount loaded | Excess GTP | Time and temperature of loading | Storage Buffer | Days from loading | GTP | GDP | GMP | % GTP loaded | Change from Day 0 |
|---|---|---|---|---|---|---|---|---|---|---|---|
| 1 | - | 0.025 mM | 10x | 10 min 37˚C | 20 mM Tris pH 8, 150 mM NaCl, 10 mM EDTA | 0 | 4.1 | 3.4 | | 54.7 | 0 |
| 2 | - | 0.025 mM | 10x | 10 min 37˚C | 20 mM Tris pH 8, 150 mM NaCl, 10 mM EDTA | 0 | 4.3 | 2.7 | | 61.5 | 0 |
| 3 | - | 0.025 mM | 10x | 10 min 37˚C | 20 mM Tris pH 8, 150 mM NaCl, 10 mM EDTA | 0 | 3.0 | 2.5 | | 54.6 | 0 |
| 4 | - | 0.025 mM | 10x | 10 min 37˚C | 20 mM Tris pH 8, 150 mM NaCl, 10 mM EDTA | 0 | 5.3 | 3.5 | | 59.8 | 0 |
| 5 | - | 0.025 mM | 10x | 30 min 37˚C | 20 mM Tris pH 8, 150 mM NaCl, 10 mM EDTA | 0 | 6.2 | 3.3 | | 65.5 | 0 |
| 6 | - | 0.025 mM | 10x | 30 min 37˚C | 20 mM Tris pH 8, 150 mM NaCl, 10 mM EDTA | 0 | 5.2 | 2.6 | | 66.6 | 0 |
| 7 | - | 0.025 mM | 10x | 30 min 37˚C | 20 mM Tris pH 8, 150 mM NaCl, 10 mM EDTA | 0 | 5.6 | 3.2 | | 64.2 | 0 |
| 8 | - | 0.025 mM | 10x | 30 min 37˚C | 20 mM Tris pH 8, 150 mM NaCl, 10 mM EDTA | 0 | 4.9 | 2.6 | | 65.7 | 0 |
| 9 | - | 0.025 mM | 10x | 1 hr 37˚C | 20 mM Tris pH 8, 150 mM NaCl, 10 mM EDTA | 0 | 5.2 | 3.0 | | 63.3 | 0 |
| 10 | - | 0.025 mM | 10x | 1 hr 37˚C | 20 mM Tris pH 8, 150 mM NaCl, 10 mM EDTA | 0 | 5.6 | 3.1 | | 64.6 | 0 |
| 11 | - | 0.025 mM | 10x | 1 hr 37˚C | 20 mM Tris pH 8, 150 mM NaCl, 10 mM EDTA | 0 | 5.3 | 2.9 | | 64.2 | 0 |
| 12 | - | 0.025 mM | 10x | 1 hr 37˚C | 20 mM Tris pH 8, 150 mM NaCl, 10 mM EDTA | 0 | 4.9 | 3.5 | | 58.2 | 0 |
| 13 | - | 0.025 mM | 10x | 1 hr RT | 20 mM Tris pH 8, 150 mM NaCl, 10 mM EDTA | 0 | 4.9 | 2.4 | | 67.5 | 0 |
| 14 | - | 0.025 mM | 10x | 1 hr RT | 20 mM Tris pH 8, 150 mM NaCl, 10 mM EDTA | 0 | 4.6 | 1.8 | | 72.3 | 0 |
| 15 | - | 0.025 mM | 10x | 1 hr RT | 20 mM Tris pH 8, 150 mM NaCl, 10 mM EDTA | 0 | 5.0 | 2.7 | | 65.2 | 0 |
| 16 | - | 0.025 mM | 10x | 1 hr RT | 20 mM Tris pH 8, 150 mM NaCl, 10 mM EDTA | 0 | 4.4 | 3.1 | | 59.2 | 0 |

| | | | | | | | | | | | |
|---|---|---|---|---|---|---|---|---|---|---|---|
| 17 | - | 0.025 mM | 10x | 1 hr 0˚C | 20 mM Tris pH 8, 150 mM NaCl, 10 mM EDTA | 0 | 5.2 | 3.0 | | 63.2 | 0 |
| 18 | - | 0.025 mM | 10x | 1 hr 0˚C | 20 mM Tris pH 8, 150 mM NaCl, 10 mM EDTA | 0 | 5.3 | 3.0 | | 64.2 | 0 |
| 19 | - | 0.025 mM | 10x | 1 hr 0˚C | 20 mM Tris pH 8, 150 mM NaCl, 10 mM EDTA | 0 | 5.5 | 3.0 | | 64.6 | 0 |
| 20 | - | 0.025 mM | 10x | 1 hr 0˚C | 20 mM Tris pH 8, 150 mM NaCl, 10 mM EDTA | 0 | 4.1 | 2.9 | | 58.2 | 0 |
| 21 | - | 0.26 mM | 10x | 1 hr RT | 20 mM Tris pH 8, 150 mM NaCl, 10 mM EDTA | 0 | 13.7 | 4.0 | 2.4 | 68.2 | 0 |
| 22 | - | 0.26 mM | 10x | 1 hr RT | 20 mM Tris pH 8, 150 mM NaCl, 10 mM EDTA | 0 | 11.4 | 5.0 | 2.4 | 60.6 | 0 |
| 23 | - | 0.26 mM | 10x | 1 hr RT | 20 mM Tris pH 8, 150 mM NaCl, 10 mM EDTA | 0 | 13.4 | 3.6 | | 78.8 | 0 |
| 24 | - | 0.26 mM | 10x | 1 hr RT | 20 mM Tris pH 8, 150 mM NaCl, 10 mM EDTA | 0 | 9.2 | 5.3 | 0.07 | 63.1 | 0 |
| 25 | - | 0.025 mM | 10x | 1 hr RT | 20 mM Tris pH 8, 150 mM NaCl, 10 mM EDTA | 0 | 11.2 | 9.1 | | 55.2 | 0 |
| 26 | + | 0.025 mM | 10x | 1 hr RT | 20 mM Tris pH 8, 150 mM NaCl, 10 mM EDTA | 0 | 5.2 | 1.0 | | 83.3 | 0 |
| 27 | + | 0.025 mM | 10x | 1 hr RT | 20 mM Tris pH 8, 150 mM NaCl, 10 mM EDTA | 0 | 5.6 | 1.2 | | 82.6 | 0 |
| 28 | + | 0.025 mM | 10x | 1 hr RT | 20 mM Tris pH 8, 150 mM NaCl, 10 mM EDTA | 0 | 5.4 | 1.6 | | 77.4 | 0 |
| 29 | + | 0.025 mM | 10x | 1 hr RT | 20 mM Tris pH 8, 150 mM NaCl, 10 mM EDTA | 0 | 4.7 | 1.9 | | 71.1 | 0 |
| 30 | + | 0.025 mM | 10x | 1 hr RT | 20 mM Tris pH 8, 150 mM NaCl, 10 mM EDTA | 0 | 11.3 | 6.8 | | 62.3 | 0 |
| 31 | - | 0.025 mM | 10x | 1 hr RT | 20 mM Tris pH 8, 150 mM NaCl, 10 mM EDTA | 0 | 11.2 | 9.1 | | 55.2 | 0 |
| 32 | - | 0.025 mM | 100x | 10 min 37˚C | 20 mM Tris pH 8, 150 mM NaCl, 10 mM EDTA | 0 | 11.3 | 2.0 | | 85.0 | 0 |
| 33 | - | 0.025 mM | 100x | 10 min 37˚C | 20 mM Tris pH 8, 150 mM NaCl, 10 mM EDTA | 0 | 12.9 | 3.2 | | 80.2 | 0 |
| 34 | - | 0.025 mM | 100x | 10 min 37˚C | 20 mM Tris pH 8, 150 mM NaCl, 10 mM EDTA | 0 | 8.3 | 2.1 | | 79.7 | 0 |
| 35 | - | 0.025 mM | 100x | 10 min 37˚C | 20 mM Tris pH 8, 150 mM NaCl, 10 mM EDTA | 0 | 8.1 | 2.1 | | 79.3 | 0 |
| 36 | - | 0.025 mM | 2.4x | 10 min 37˚C | 20 mM Tris pH 8, 150 mM NaCl, | 0 | 4.1 | 4.9 | | 45.6 | 0 |

| | | | | | | | | | | | |
|---|---|---|---|---|---|---|---|---|---|---|---|
| | | | | | 10 mM EDTA | | | | | | |
| 37 | - | 0.025 mM | 2.4x | 10 min 37˚C | 20 mM Tris pH 8, 150 mM NaCl, 10 mM EDTA | 0 | 4.3 | 5.1 | | 45.8 | 0 |
| 38 | - | 0.025 mM | 2.4x | 10 min 37˚C | 20 mM Tris pH 8, 150 mM NaCl, 10 mM EDTA | 0 | 3.7 | 5.8 | | 38.9 | 0 |
| 39 | - | 0.025 mM | 2.4x | 10 min 37˚C | 20 mM Tris pH 8, 150 mM NaCl, 10 mM EDTA | 0 | 4.8 | 7.1 | | 39.9 | 0 |
| 40 | + | 0.025 mM | 100x | 10 min 37˚C | 20 mM Tris pH 8, 150 mM NaCl, 10 mM EDTA | 0 | 3.4 | 1.3 | | 73.2 | 0 |
| 41 | + | 0.025 mM | 100x | 10 min 37˚C | 20 mM Tris pH 8, 150 mM NaCl, 10 mM EDTA | 0 | 7.0 | 2.1 | | 77.0 | 0 |
| 42 | + | 0.025 mM | 100x | 10 min 37˚C | 20 mM Tris pH 8, 150 mM NaCl, 10 mM EDTA | 0 | 5.9 | 2.9 | | 67.8 | 0 |
| 43 | + | 0.025 mM | 100x | 10 min 37˚C | 20 mM Tris pH 8, 150 mM NaCl, 10 mM EDTA | 0 | 5.9 | 2.1 | | 73.7 | 0 |
| 44 | - | 0.025 mM | 10x | 1 hr 0˚C | 20 mM Tris pH 8, 150 mM NaCl, 10 mM $MgCl_2$ | 0 | 3.6 | 4.2 | | 46.2 | 0 |
| 45 | - | 0.025 mM | 10x | 1 hr 0˚C | 20 mM Tris pH 8, 150 mM NaCl, 10 mM $MgCl_2$ | 0 | 4.0 | 3.0 | | 57.1 | 0 |
| 46 | - | 0.025 mM | 10x | 1 hr 0˚C | 20 mM Tris pH 8, 150 mM NaCl, 10 mM $MgCl_2$ | 0 | 3.0 | 4.6 | | 39.3 | 0 |
| 47 | - | 0.025 mM | 10x | 1 hr 0˚C | 20 mM Tris pH 8, 150 mM NaCl, 10 mM $MgCl_2$ | 0 | 5.2 | 3.7 | | 58.1 | 0 |
| 48 | - | 0.025 mM | 10x | 1 hr 0˚C | 20 mM Tris pH 8, 150 mM NaCl | 0 | 5.5 | 3.2 | | 63.1 | 0 |
| 49 | - | 0.025 mM | 10x | 1 hr 0˚C | 20 mM Tris pH 8, 150 mM NaCl | 0 | 6.0 | 3.4 | | 64.1 | 0 |
| 50 | - | 0.025 mM | 10x | 1 hr 0˚C | 20 mM Tris pH 8, 150 mM NaCl | 0 | 6.2 | 3.5 | | 64.1 | 0 |
| 51 | - | 0.025 mM | 10x | 1 hr 0˚C | 20 mM Tris pH 8, 150 mM NaCl | 0 | 5.0 | 3.2 | | 61.1 | 0 |
| 52 | - | 0.025 mM | 10x | 1 hr 0˚C | 20 mM Tris pH 8, 150 mM NaCl, 10 mM EDTA, 10 mM $MgCl_2$ | 0 | 2.7 | 4.6 | | 36.8 | 0 |
| 53 | - | 0.025 mM | 10x | 1 hr 0˚C | 20 mM Tris pH 8, 150 mM NaCl, 10 mM EDTA, 10 mM $MgCl_2$ | 0 | 2.6 | 4.8 | | 35.0 | 0 |
| 54 | - | 0.025 mM | 10x | 1 hr 0˚C | 20 mM Tris pH 8, 150 mM NaCl, 10 mM EDTA, 10 mM $MgCl_2$ | 0 | 2.7 | 5.1 | | 34.8 | 0 |
| 55 | - | 0.025 mM | 10x | 1 hr 0˚C | 20 mM Tris pH 8, 150 mM NaCl, 10 mM EDTA, 10 mM $MgCl_2$ | 0 | 2.7 | 5.0 | | 35.4 | 0 |
| 56 | - | 0.025 mM | 10x | 1 hr 0˚C | 20 mM Tris pH 8, 150 mM NaCl | 0 | 11.0 | 7.3 | | 60.2 | 0.0 |

| | | | | | | | | | | | |
|---|---|---|---|---|---|---|---|---|---|---|---|
| 57 | - | 0.025 mM | 10x | 1 hr 0˚C | 20 mM Tris pH 8, 150 mM NaCl | 1 | 6.4 | 11.6 | | 35.6 | -24.5 |
| 58 | - | 0.025 mM | 10x | 1 hr 0˚C | 20 mM Tris pH 8, 150 mM NaCl | 3 | 0.1 | 2.0 | | 6.5 | -53.6 |
| 59 | - | 0.025 mM | 10x | 10 min 37˚C | 20 mM Tris pH 8, 150 mM NaCl | 1 | 6.3 | 3.3 | | 65.2 | -19.9 |
| 60 | - | 0.025 mM | 10x | 10 min 37˚C | 20 mM Tris pH 8, 150 mM NaCl | 3 | 1.8 | 5.1 | | 26.1 | -58.9 |
| 61 | - | 0.025 mM | 10x | 10 min 37˚C | 20 mM Tris pH 8, 150 mM NaCl | 1 | 11.5 | 3.9 | | 74.4 | -5.8 |
| 62 | - | 0.025 mM | 10x | 10 min 37˚C | 20 mM Tris pH 8, 150 mM NaCl | 3 | 7.4 | 5.2 | | 58.6 | -21.6 |
| 63 | - | 0.025 mM | 10x | 10 min 37˚C | 20 mM Tris pH 8, 150 mM NaCl | 0 | 3.4 | 3.3 | | 50.8 | 0.0 |
| 64 | - | 0.025 mM | 10x | 10 min 37˚C | 20 mM Tris pH 8, 150 mM NaCl | 1 | 2.7 | 4.2 | | 39.4 | -11.4 |
| 65 | - | 0.025 mM | 10x | 10 min 37˚C | 20 mM Tris pH 8, 150 mM NaCl | 3 | 1.8 | 5.1 | | 26.1 | -24.7 |
| 66 | - | 0.025 mM | 10x | 10 min 37˚C | 20 mM Tris pH 8, 150 mM NaCl, 10 mM EDTA | 1 | 4.4 | 3.6 | | 55.0 | 0.3 |
| 67 | - | 0.025 mM | 10x | 10 min 37˚C | 20 mM Tris pH 8, 150 mM NaCl, 10 mM EDTA | 3 | 3.8 | 3.5 | | 52.1 | -2.7 |
| 68 | - | 0.025 mM | 10x | 10 min 37˚C | 20 mM Tris pH 8, 150 mM NaCl, 10 mM EDTA | 1 | 3.0 | 2.4 | | 55.9 | -5.5 |
| 69 | - | 0.025 mM | 10x | 10 min 37˚C | 20 mM Tris pH 8, 150 mM NaCl, 10 mM EDTA | 3 | 2.5 | 2.2 | | 53.8 | -7.7 |
| 70 | - | 0.025 mM | 10x | 10 min 37˚C | 20 mM Tris pH 8, 150 mM NaCl, 10 mM EDTA | 1 | 2.9 | 2.4 | | 54.3 | -0.3 |
| 71 | - | 0.025 mM | 10x | 10 min 37˚C | 20 mM Tris pH 8, 150 mM NaCl, 10 mM EDTA | 3 | 2.3 | 2.1 | | 52.3 | -2.3 |
| 72 | - | 0.025 mM | 10x | 10 min 37˚C | 20 mM Tris pH 8, 150 mM NaCl, 10 mM EDTA | 0 | 4.6 | 2.9 | | 61.0 | 0.0 |
| 73 | - | 0.025 mM | 10x | 10 min 37˚C | 20 mM Tris pH 8, 150 mM NaCl, 10 mM EDTA | 1 | 4.0 | 3.2 | | 55.1 | -5.9 |
| 74 | - | 0.025 mM | 10x | 10 min 37˚C | 20 mM Tris pH 8, 150 mM NaCl, 10 mM EDTA | 3 | 3.3 | 3.2 | | 50.6 | -10.4 |
| 75 | - | 0.025 mM | 10x | 1 hr 0˚C | 20 mM Tris pH 8, 150 mM NaCl, 10 mM $MgCl_2$ | 1 | 1.9 | 4.7 | | 28.3 | -17.8 |
| 76 | - | 0.025 mM | 10x | 1 hr 0˚C | 20 mM Tris pH 8, 150 mM NaCl, 10 mM $MgCl_2$ | 3 | 0.8 | 4.9 | | 13.6 | -32.6 |
| 77 | - | 0.025 mM | 10x | 10 min 37˚C | 20 mM Tris pH 8, 150 mM NaCl, 10 mM $MgCl_2$ | 0 | 3.2 | 4.0 | | 44.7 | 0.0 |
| 78 | - | 0.025 mM | 10x | 10 min 37˚C | 20 mM Tris pH 8, 150 mM NaCl, 10 mM $MgCl_2$ | 1 | 1.8 | 5.1 | | 25.5 | -19.2 |

| | | | | | | | | | | | |
|---|---|---|---|---|---|---|---|---|---|---|---|
| 79 | - | 0.025 mM | 10x | 10 min 37˚C | 20 mM Tris pH 8, 150 mM NaCl, 10 mM $MgCl_2$ | 3 | 0.9 | 5.4 | | 14.6 | -30.1 |
| 80 | - | 0.025 mM | 10x | 10 min 37˚C | 20 mM Tris pH 8, 150 mM NaCl, 10 mM $MgCl_2$ | 0 | 4.6 | 3.1 | | 59.5 | 0.0 |
| 81 | - | 0.025 mM | 10x | 10 min 37˚C | 20 mM Tris pH 8, 150 mM NaCl, 10 mM $MgCl_2$ | 1 | 2.5 | 4.4 | | 36.4 | -23.0 |
| 82 | - | 0.025 mM | 10x | 10 min 37˚C | 20 mM Tris pH 8, 150 mM NaCl, 10 mM $MgCl_2$ | 3 | 1.3 | 3.9 | | 24.5 | -34.9 |
| 83 | - | 0.025 mM | 10x | 10 min 37˚C | 20 mM Tris pH 8, 150 mM NaCl, 10 mM $MgCl_2$ | 0 | 3.9 | 2.7 | | 59.1 | 0.0 |
| 84 | - | 0.025 mM | 10x | 10 min 37˚C | 20 mM Tris pH 8, 150 mM NaCl, 10 mM $MgCl_2$ | 1 | 2.3 | 5.0 | | 31.4 | -27.7 |
| 85 | - | 0.025 mM | 10x | 10 min 37˚C | 20 mM Tris pH 8, 150 mM NaCl, 10 mM $MgCl_2$ | 3 | 1.5 | 5.1 | | 22.9 | -36.3 |
| 86 | - | 0.025 mM | 10x | 1 hr 0˚C | 20 mM Tris pH 8, 150 mM NaCl, 10 mM EDTA, 10 mM $MgCl_2$ | 0 | 3.0 | 3.6 | | 45.2 | 0.0 |
| 87 | - | 0.025 mM | 10x | 1 hr 0˚C | 20 mM Tris pH 8, 150 mM NaCl, 10 mM EDTA, 10 mM $MgCl_2$ | 1 | 2.9 | 4.2 | | 40.7 | -4.5 |
| 88 | - | 0.025 mM | 10x | 1 hr 0˚C | 20 mM Tris pH 8, 150 mM NaCl, 10 mM EDTA, 10 mM $MgCl_2$ | 3 | 1.0 | 5.2 | | 16.2 | -28.9 |
| 89 | - | 0.025 mM | 10x | 10 min 37˚C | 20 mM Tris pH 8, 150 mM NaCl, 10 mM EDTA, 10 mM $MgCl_2$ | 0 | 3.1 | 3.8 | | 45.1 | 0.0 |
| 90 | - | 0.025 mM | 10x | 10 min 37˚C | 20 mM Tris pH 8, 150 mM NaCl, 10 mM EDTA, 10 mM $MgCl_2$ | 1 | 2.1 | 4.8 | | 30.3 | -14.8 |
| 91 | - | 0.025 mM | 10x | 10 min 37˚C | 20 mM Tris pH 8, 150 mM NaCl, 10 mM EDTA, 10 mM $MgCl_2$ | 3 | 0.9 | 5.4 | | 14.7 | -30.3 |
| 92 | - | 0.025 mM | 10x | 10 min 37˚C | 20 mM Tris pH 8, 150 mM NaCl, 10 mM EDTA, 10 mM $MgCl_2$ | 0 | 2.7 | 3.5 | | 43.1 | 0.0 |
| 93 | - | 0.025 mM | 10x | 10 min 37˚C | 20 mM Tris pH 8, 150 mM NaCl, 10 mM EDTA, 10 mM $MgCl_2$ | 1 | 2.0 | 4.4 | | 31.0 | -12.1 |
| 94 | - | 0.025 mM | 10x | 10 min 37˚C | 20 mM Tris pH 8, 150 mM NaCl, 10 mM EDTA, 10 mM $MgCl_2$ | 3 | 0.9 | 5.3 | | 14.6 | -28.5 |
| 95 | - | 0.025 mM | 10x | 10 min 37˚C | 20 mM Tris pH 8, 150 mM NaCl, 10 mM EDTA, 10 mM $MgCl_2$ | 0 | 3.7 | 4.4 | | 45.7 | 0.0 |
| 96 | - | 0.025 mM | 10x | 10 min 37˚C | 20 mM Tris pH 8, 150 mM NaCl, 10 mM EDTA, 10 mM $MgCl_2$ | 1 | 2.4 | 4.7 | | 33.6 | -12.2 |
| 97 | - | 0.025 mM | 10x | 10 min 37˚C | 20 mM Tris pH 8, 150 mM NaCl, 10 mM EDTA, 10 mM $MgCl_2$ | 3 | 1.2 | 4.6 | | 20.7 | -25.1 |

**Supplemental Table 2. The conditions and replicates of the nucleotide content of the loaded K-Ras and N-Ras isoforms.**

| **Loaded Ras Isoforms** | | | | | | | | | | |
|---|---|---|---|---|---|---|---|---|---|---|
| **Trial #** | **Mg2+ Addition** | **Amount Loaded** | **Excess GTP** | **Time and temperature of loading** | **Storage Buffer** | **Isoform** | **GTP** | **GDP** | **GMP** | **% GTP loaded** |
| 98 | - | 0.025 mM | 10x | 10 min 37˚C | 20 mM Tris pH 8, 150 mM NaCl, 10 mM EDTA | K-Ras | **3.8** | **2.8** | | **57.1** |
| 99 | - | 0.025 mM | 10x | 10 min 37˚C | 20 mM Tris pH 8, 150 mM NaCl, 10 mM EDTA | K-Ras | 4.1 | 3.9 | | 51.3 |
| 100 | - | 0.025 mM | 10x | 10 min 37˚C | 20 mM Tris pH 8, 150 mM NaCl, 10 mM EDTA | K-Ras | 3.4 | 3.3 | | 51.3 |
| 101 | - | 0.025 mM | 10x | 10 min 37˚C | 20 mM Tris pH 8, 150 mM NaCl, 10 mM EDTA | K-Ras | 3.2 | 2.8 | | 53.4 |
| 102 | - | 0.025 mM | 10x | 10 min 37˚C | 20 mM Tris pH 8, 150 mM NaCl, 10 mM EDTA | N-Ras | 2.2 | 1.4 | | 61.1 |
| 103 | - | 0.025 mM | 10x | 10 min 37˚C | 20 mM Tris pH 8, 150 mM NaCl, 10 mM EDTA | N-Ras | 3.3 | 2.4 | | 57.7 |
| 104 | - | 0.025 mM | 10x | 10 min 37˚C | 20 mM Tris pH 8, 150 mM NaCl, 10 mM EDTA | N-Ras | 3.1 | 2.1 | | 59.5 |
| 105 | - | 0.025 mM | 10x | 10 min 37˚C | 20 mM Tris pH 8, 150 mM NaCl, 10 mM EDTA | N-Ras | 3.3 | 2.2 | | 60.2 |

**Supplemental Table 3. Nucleotide content of purified wild-type and mutant H-Ras.**

| Purified Cancer Mutations | | | | |
|---|---|---|---|---|
| **Mutation** | **GTP** | **GDP** | **GMP** | **% GTP** |
| WT | 1.7 | 16.9 | 9.8 | 6.0 |
| WT | 0.8 | 9.8 | 7.9 | 4.3 |
| WT | 2.5 | 37.5 | 13.8 | 4.6 |
| WT | 2.3 | 38.0 | 14.3 | 4.3 |
| G12V | 15.0 | 13.4 | 14.0 | 35.4 |
| G12V | 11.8 | 21.6 | 17.9 | 23.1 |
| G12V | 10.7 | 24.7 | 10.3 | 23.4 |
| G12V | 14.1 | 29.8 | 16.1 | 23.6 |
| G13R | 2.5 | 11.5 | 10.6 | 10.1 |
| G13R | 8.5 | 34.8 | 17.5 | 14.0 |
| G13R | 6.5 | 38.3 | 19.4 | 10.1 |
| G13R | 7.4 | 29.9 | 15.1 | 14.2 |
| Q61L | 21.2 | 20.2 | 26.7 | 31.1 |
| Q61L | 18.4 | 19.4 | 24.6 | 29.4 |
| Q61L | 18.1 | 23.7 | 10.2 | 34.9 |
| Q61L | 19.3 | 23.7 | 19.3 | 31.0 |

**Supplemental Table 4. The conditions and replicates of the nucleotide content of loaded H-Ras-G12V.**

| Loaded Cancer Mutations | | | | | | | | | | |
|---|---|---|---|---|---|---|---|---|---|---|
| **Trial #** | **Mg2+ Addition** | **Amount Loaded** | **Excess GTP** | **Time and temperature of loading** | **Storage Buffer** | **Mutation** | **GTP** | **GDP** | **GMP** | **% GTP loaded** |
| 106 | - | 0.025 mM | 10x | 10 min 37˚C | 20 mM Tris pH 8, 150 mM NaCl, 10 mM EDTA | G12V | 14.3 | 25.7 | | 35.8 |
| 107 | - | 0.025 mM | 10x | 10 min 37˚C | 20 mM Tris pH 8, 150 mM NaCl, 10 mM EDTA | G12V | 8.6 | 9.5 | | 47.5 |
| 108 | - | 0.025 mM | 10x | 10 min 37˚C | 20 mM Tris pH 8, 150 mM NaCl, 10 mM EDTA | G12V | 6.2 | 11.6 | | 34.8 |
| 109 | - | 0.025 mM | 10x | 10 min 37˚C | 20 mM Tris pH 8, 150 mM NaCl, 10 mM EDTA | G12V | 3.6 | 10.4 | | 25.4 |
